# MoleculeExperiment enables consistent infrastructure for molecule-resolved spatial transcriptomics data in Bioconductor

**DOI:** 10.1101/2023.05.16.541040

**Authors:** Bárbara Zita Peters Couto, Nicholas Robertson, Ellis Patrick, Shila Ghazanfar

## Abstract

Imaging-based spatial transcriptomics technologies have achieved subcellular resolution, enabling detection of individual molecules in their native tissue context. Data associated with these technologies promises unprecedented opportunity towards understanding cellular and subcellular biology. However, in R/Bioconductor there is a scarcity of existing computational infrastructure to represent such data, and particularly to summarise and transform it for existing widely adopted computational tools in single cell transcriptomics analysis, including SingleCellExperiment and SpatialExperiment classes. With the emergence of several commercial offerings of imaging-based spatial transcriptomics, there is a pressing need to develop consistent data structure standards for these technologies at the individual molecule level. To this end, we have developed MoleculeExperiment, an R/Bioconductor package, which i) stores molecule and cell segmentation boundary information at the molecule-level, ii) standardises this molecule-level information across different imaging-based ST technologies, including 10x Genomics’ Xenium, and iii) streamlines transition from a MoleculeExperiment object to a SpatialExperiment object. Overall, MoleculeExperiment is generally applicable as a data infrastructure class for consistent analysis of imaging-based spatial transcriptomics data.

## Introduction

Spatial omics is a maturing field, especially spatial transcriptomics (ST) technologies^1^. Since the publication of single-molecule FISH, many imaging-based ST technologies have been developed^2^. While the transcriptome coverage of these technologies is not complete, they enable cellular and even subcellular resolution^1^. In addition, various imaging-based ST technologies have recently started to be commercially shipped, such as 10x Genomics’s Xenium^3^, and thus their use is expected to massively increase in scale. Imaging-based ST has been employed in multiple studies, including investigating the progression from ductal carcinoma in situ to invasive carcinoma^3^, analysing the complex immune landscape of the tumour microenvironment in lung tumours^4^, and facilitating the creation of the first comprehensive spatial atlas of the mouse brain^5^. The highly informative cellular and subcellular resolution of imaging-based ST, as well as its increasing commercial availability, motivate the generation of software that helps scientists consistently handle this type of high resolution data.

Recently, SpatialExperiment (SE) was developed as an object class for the study of spatial transcriptomics data^6^. Just like the commonly used SingleCellExperiment class, SpatialExperiment aims to promote reproducibility of analyses and interoperability of different software on the same data^6,7^. Moreover, by being part of the Bioconductor project, these packages are part of an effort to disseminate open data analysis and promote software maintenance and enhancement in the life sciences^7^. However, the SE package only allows the storage of information at the cellular level. Thus, transcripts that have not been assigned to a cell by the cell segmentation method are lost, which is disadvantageous, as these transcripts could yield valuable biological insights^8^. For example, performing a region-level differential expression analysis could be more accurate if all detected transcripts are taken into account, even in spaces where transcripts have not been assigned to a cell. However, if one were interested in doing such an analysis on ST data that has been summarised as an SE object, one would only be able to do this at the cell-level. Therefore, to leverage the molecule resolution of recent technologies, there is a need for a class that avoids premature summarisation of ST data, and enables analysis of transcripts in their 3D locations irrespective of cellular compartmentalisation.

In this paper, we introduce the MoleculeExperiment class, which represents ST data at the molecule-level. In addition, the MoleculeExperiment class imposes standardised data formats and terminology to avoid the need for manual file conversion and complex analysis scripts of molecule-based ST data. Moreover, the MoleculeExperiment package facilitates the transition to a cell-level analysis with the already existing SpatialExperiment class. Here we primarily focus on the application of MoleculeExperiment to Xenium (10x Genomics), CosMx (NanoString) and MERSCOPE (Vizgen) data. In summary, the MoleculeExperiment package aims to facilitate the downstream analysis of different imaging-based ST data, both at the molecule-level and cellular-level, with the large diversity of data analysis tools in the Bioconductor project.

## Results

Here we introduce MoleculeExperiment, a core data infrastructure package in R/Bioconductor which enables consistent and reproducible analysis of molecule resolution spatial transcriptomics data in the R coding environment (Figure 1). The `MoleculeExperiment` class is an S4 class with two primary slots for storing information on `molecules` and/or `boundaries`. The `molecules` slot stores information about the detected molecules and is nested by `assay`, e.g. for different transcript decoding approaches^9,10^, by `sample` for datasets with multiple samples and/or images, and by `feature_name` for different transcripts or molecules. The core information in `feature_name` is the `x_location` and `y_location` of each molecule, but other information can be stored here. The `boundaries` slot is used for storing various segmentations of the data and is also nested by `assay`, for different segmentations such as cell bodies, nuclei or tissue compartment, by `sample`, and by `segment_id` for each individual segment (typically a cell).

**Figure 1.**
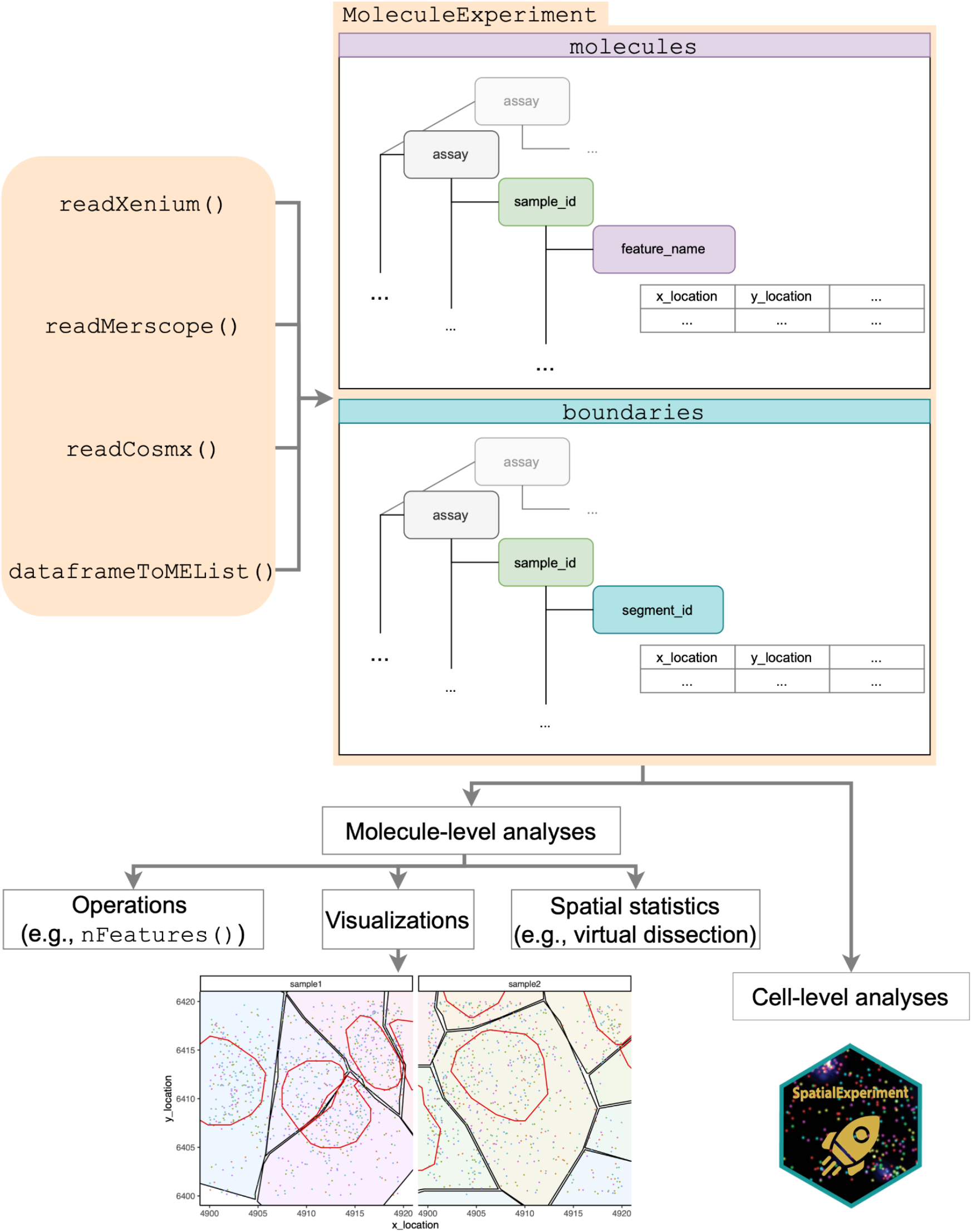
MoleculeExperiment aims to facilitate molecule-level and cell-level analysis of data across different vendors of imaging-based ST data. A `MoleculeExperiment` object has a `molecules` slot and a `boundaries` slot, where data format and terminology are standardised (e.g., hierarchical nested list for storage, µm units for the coordinates, and specific column names). The `MoleculeExperiment` class enables a molecule-centric analysis via class-specific accessor functions. Possible molecule-level downstream analyses include visualisations (e.g., digital *in situs*), operations (e.g., filtering and counting), and spatial statistics (e.g., Differential gene expression (DGE) by virtual dissection). In addition, the MoleculeExperiment package facilitates transition to a cell level analysis via the summarisation of molecule-level data into a SpatialExperiment object.

Due to the large variation in data bundles produced by various vendors (Supplementary Figure 1), we have implemented specific functions for reading and standardising this data into a `MoleculeExperiment` object. Currently, we have implemented are `readXenium()`, `readCosmx()` and `readMerscope(),` alongside a technology-agnostic `dataFrameToMEList()` function. The package provides setter and getter functions, e.g., `molecules()` and `boundaries()`, needed to manipulate the object in R.

The hierarchical nested structure of the `MoleculeExperiment` class avoids redundant storing of information, as opposed to traditional rectangular data storage formats (e.g., csv files). For example, the sample IDs and feature names are not repeated for the millions of molecules corresponding to that sample and feature. As such, `MoleculeExperiment` creates objects that consume less memory than rectangular objects (Supplementary Figure 2). Avoiding redundancy is of special interest when working with such high-throughput technologies.

The molecule-centric way in which the `MoleculeExperiment` object stores data can be used for molecule-level visualisations and statistical analyses (Figure 1). Further, we provide the `countMolecules()` function to count the number of molecules in each segment and produce a `SpatialExperiment` object. Thus, the `MoleculeExperiment` object not only takes advantage of the molecule resolution of imaging-based ST technologies, but also facilitates the transition from a molecule-level analysis to a cell-level analysis, thereby leveraging the vast capacity of Bioconductor tools.

## Discussion

Here, we have developed MoleculeExperiment, an S4 infrastructure in R/Bioconductor, for molecule resolved omics data. First, the MoleculeExperiment class enables analysis of imaging-based spatial transcriptomics data at the molecule level, thereby making full use of the molecule resolution that these technologies can achieve. Second, it imposes a standardisation of the data such that the data structure, and associated terminology, are consistent across data from diverse vendors of imaging-based ST. This consistent data representation aims to provide a solid foundation for the development of tools for analysis of molecule-level ST data. We think this is especially important in the current context, where new imaging-based ST technologies and associated analytical methods are constantly being developed^1,2^. Third, the MoleculeExperiment package provides convenience functions to read in data from the different vendors of imaging-based ST technologies. This aims to simplify the otherwise manual and time-consuming process of in-house data handling and reformatting before data analyses. Lastly, the package facilitates the transition into a SpatialExperiment object for cell-level analyses. In this way, it is possible to use already existing Bioconductor packages that work with the SpatialExperiment or SingleCellExperiment classes, e.g., imcRtools^11^, spicyR^12^, SPIAT^13^ and scHOT^14^. Moreover, this means that the molecule-resolved data can be transitioned to other related classes, like SingleCellExperiment^7^ and its python-equivalent AnnData^15^. Taken together, by being an S4 Bioconductor class, MoleculeExperiment profits from interoperability of downstream software packages, like the SpatialExperiment^6^ and SingleCellExperiment^7^ classes do. In summary, the MoleculeExperiment package i) enables molecule-level analyses, ii) standardises data formats across different vendors, and iii) facilitates transition from a MoleculeExperiment object to a SpatialExperiment object for cell-level analyses with existing tools in the Bioconductor project. Ultimately, the MoleculeExperiment package imposes a consistent structure and terminology for imaging-based ST data, with the goal of enabling reproducible downstream molecular- and cellular-level analysis for the user.

A key advantage of the nested structure of MoleculeExperiment is that it facilitates parallelisation and on-disk handling of data. The hierarchical nested format of the MoleculeExperiment data enables parallelisation via BiocParallel^16^ and may be handled on-disk via tools such as the Apache Arrow package (https://arrow.apache.org/) or HDF5^17^ as is done in other Bioconductor packages^18^. While current molecule-level ST have few replicates, as these technologies increase to cohort-scale experiments, the capacity to not retain the whole dataset in memory for analysis will be vital.

In summary, the MoleculeExperiment R package standardises imaging-based ST data at the molecule level across different vendors, and simplifies the steps needed to prepare raw imaging-based ST data, ready for downstream analyses at the cellular and molecular level. We hope MoleculeExperiment supports the recent and fast-growing spatial omics community.

### Availability and Implementation

The MoleculeExperiment package is publicly available on Bioconductor at https://bioconductor.org/packages/release/bioc/html/MoleculeExperiment.html. Source code is available on Github at: https://github.com/SydneyBioX/MoleculeExperiment. The vignette for MoleculeExperiment can be found at https://bioconductor.org/packages/release/bioc/html/MoleculeExperiment.html.

## Methods

### Examination of vendors’ public molecule-resolved spatial transcriptomics data structure

We examined molecule and boundary data structures from the following technologies: 10x Genomics Xenium, NanoString CosMx, and Vizgen MERSCOPE. We used these vendors’ publicly available output data bundles, in some cases requiring a minimal sign in or form completion. We used these data bundles to inform our `readXenium`, `readCosmx` and `readMerscope` functions respectively. In particular, Xenium data corresponds to fresh frozen mouse brain tissue (https://www.10xgenomics.com/resources/datasets/fresh-frozen-mouse-brain-replicates-1-standard), accessed on 8 February 2023; CosMx data corresponds to human non-small cell lung cancer (https://nanostring.com/resources/smi-ffpe-dataset-lung9-rep1-data/) accessed on 27 February 2023; and MERSCOPE data is from human ovarian cancer (https://console.cloud.google.com/storage/browser/vz-ffpe-showcase/HumanOvarianCancerPatient2Slice2) accessed on 27 February 2023.

We assessed commonalities in terms of the detected transcripts files as well as cell boundary or segmentation files. No commonalities were found in the cell boundary files across the technologies. Vizgen’s output bundle contains several hdf5 files, Xenium a single csv.gz file, and NanoString has no single file with cell boundaries, but shares the identified cell IDs between the transcript, count matrix, and cell metadata files instead.

### Assessing MoleculeExperiment memory

To assess the disk and memory size of molecule data objects, we used the public CosMx data corresponding to the “Lung9_Rep1” sample, with 26,275,891 molecules detected over 900 features. We assessed on disk file sizes for the transcript csv file as made available from the NanoString website, a Gzip compressed csv.gz version of the file, as well as MoleculeExperiment objects exported to disk via readRDS, either including all additional columns or only keeping essential columns. To assess memory sizes, we compared the two MoleculeExperiment objects to a data.frame generated by reading the aforementioned csv file. We quantified file and object sizes using the file.size and object.size functions respectively and reported these in megabytes (MB).

## BACKMATTER

### Ethics statement

Not applicable.

## Acknowledgments

The authors thank all their colleagues, particularly at The University of Sydney, Sydney Precision Data Science and Judith and David Coffey Life Lab in Charles Perkins Centre for their support and intellectual engagement. We especially thank Nils Eling and Ludwig Geistlinger, along with all members of the Spatial Imaging Special Interest Group, for their careful feedback and engagement.

## Author Contribution

SG, EP, conceived, designed and funded the study. BZPC completed the analysis and design of software with feedback from SG and EP. BZPC and NR implemented and constructed the R package with feedback from SG and EP. BZPC and NR tested the R package; All authors wrote, reviewed and approved the manuscript.

## Supplementary Materials

### Supplementary Figures

**Supplementary Figure 1.**
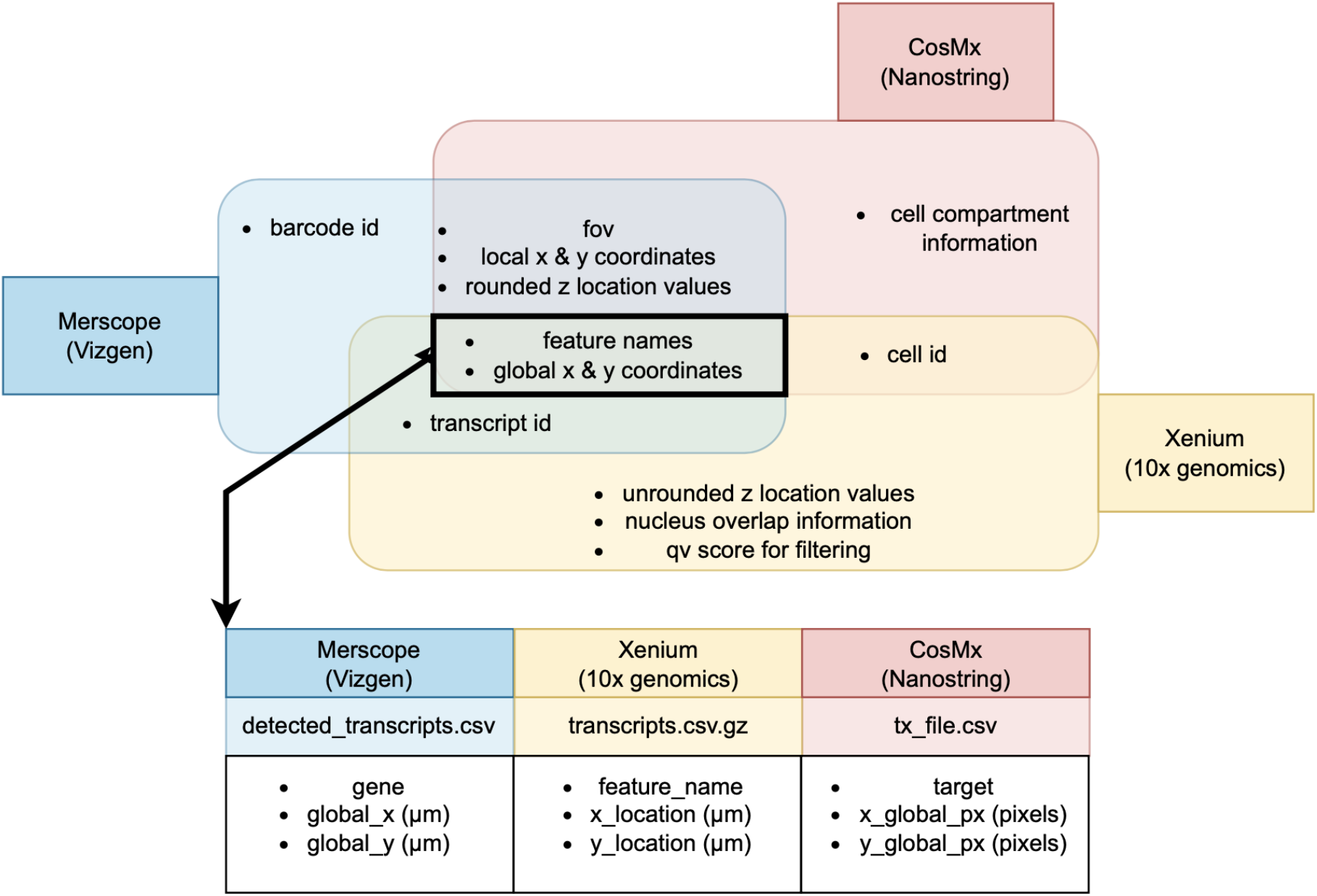
Assessment of consistency of data structure among different imaging-based spatial transcriptomics vendors. Venn diagram showing label and type of information in the detected transcript files among technology vendors. Only i) the feature names (including gene names), and ii) the global x and y coordinates of those features are shared. Common information displays different file compression formats, column units and names.

**Supplementary Figure 2.**
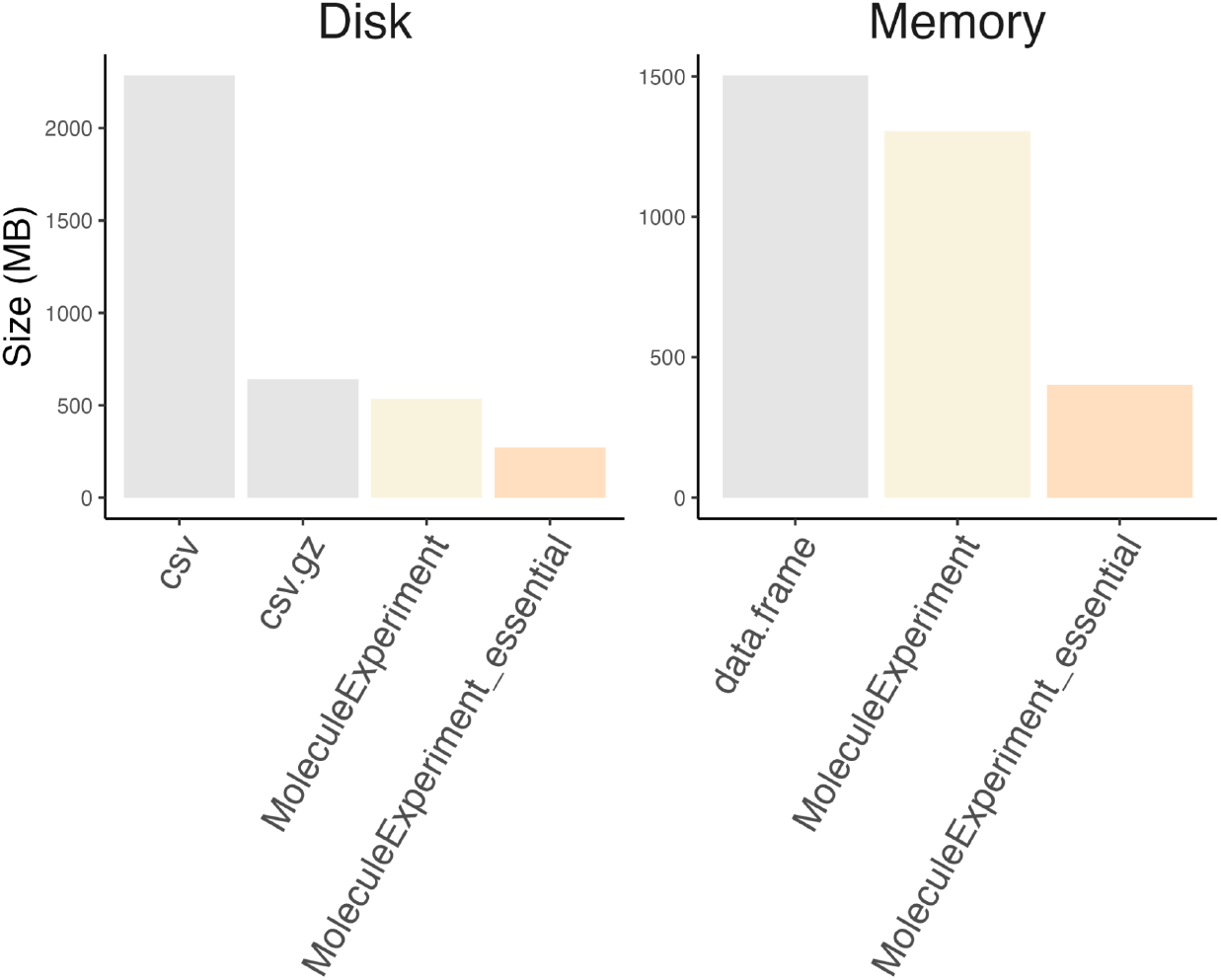
Relative memory use of MoleculeExperiment. Barplot displaying memory usage on disk or in memory (MB) of NanoString CosMx molecule-level data for publicly available sample “Lung9_Rep1” when using various infrastructure approaches.

## References

1. Wu, Y., Cheng, Y., Wang, X., Fan, J. & Gao, Q. Spatial omics: Navigating to the golden era of cancer research. Clin. Transl. Med. 12, e696 (2022).

2. Williams, C. G., Lee, H. J., Asatsuma, T., Vento-Tormo, R. & Haque, A. An introduction to spatial transcriptomics for biomedical research. Genome Med. 14, 68 (2022).

3. Janesick, A. et al. High resolution mapping of the breast cancer tumor microenvironment using integrated single cell, spatial and in situ analysis of FFPE tissue. bioRxiv 2022.10.06.510405 (2022) doi:10.1101/2022.10.06.510405.

4. Chen, J. H. et al. Spatial analysis of human lung cancer reveals organized immune hubs enriched for stem-like CD8 T cells and associated with immunotherapy response. bioRxiv 2023.04.04.535379 (2023) doi:10.1101/2023.04.04.535379.

5. Yao, Z. et al. A high-resolution transcriptomic and spatial atlas of cell types in the whole mouse brain. bioRxiv 2023.03.06.531121 (2023) doi:10.1101/2023.03.06.531121.

6. Righelli, D. et al. SpatialExperiment: infrastructure for spatially-resolved transcriptomics data in R using Bioconductor. Bioinformatics 38, 3128–3131 (2022).

7. Amezquita, R. A. et al. Orchestrating single-cell analysis with Bioconductor. Nat. Methods 17, 137–145 (2020).

8. Prabhakaran, S. Sparcle: assigning transcripts to cells in multiplexed images. Bioinform Adv 2, vbac048 (2022).

9. Gataric, M. et al. PoSTcode: Probabilistic image-based spatial transcriptomics decoder. bioRxiv 2021.10.12.464086 (2021) doi:10.1101/2021.10.12.464086.

10. Cisar, C., Keener, N., Ruffalo, M. & Paten, B. A Unified Pipeline for FISH Spatial Transcriptomics. bioRxiv 2023.02.17.529010 (2023) doi:10.1101/2023.02.17.529010.

11. Windhager, J., Bodenmiller, B. & Eling, N. An end-to-end workflow for multiplexed image processing and analysis. bioRxiv 2021.11.12.468357 (2021) doi:10.1101/2021.11.12.468357.

12. Canete, N.. et al. spicyR: spatial analysis of in situ cytometry data in R. Bioinformatics 38, 3099–3105 (2022).

13. Yang, T. et al. SPIAT: An R package for the Spatial Image Analysis of Cells in Tissues. bioRxiv 2020.05.28.122614 (2020) doi:10.1101/2020.05.28.122614.

14. Ghazanfar, S. et al. Investigating higher-order interactions in single-cell data with scHOT. Nature (2020).

15. Virshup, I., Rybakov, S., Theis, F. J., Angerer, P. & Alexander Wolf, F. anndata: Annotated data. bioRxiv 2021.12.16.473007 (2021) doi:10.1101/2021.12.16.473007.

16. Morgan, M., Obenchain, V., Lang, M., Thompson, R. & Turaga, N. BiocParallel: Bioconductor facilities for parallel evaluation. 2016. URL https://bioconductor.org/packages/BiocParallel. R package version 1, (2017).

17. Fischer, B., Pau, G. & Smith, M. rhdf5: HDF5 interface to R. R package version 2, (2017).

18. Eling, N., Damond, N., Hoch, T. & Bodenmiller, B. cytomapper: an R/Bioconductor package for visualization of highly multiplexed imaging data. Bioinformatics 36, 5706–5708 (2021).

19. Dries, R. et al. Advances in spatial transcriptomic data analysis. Genome Res. 31, 1706–1718 (2021).

